# Ankylosis homologue (ANKH) controls extracellular citrate and pyrophosphate homeostasis and affects bone mechanical performance

**DOI:** 10.1101/2019.12.20.883223

**Authors:** Flora Szeri, Stefan Lundkvist, Sylvia Donnelly, Udo F.H. Engelke, Kyu Rhee, Charlene Williams, John P. Sundberg, Ron A. Wevers, Ryan E. Tomlinson, Robert Jansen, Koen van de Wetering

**Author notes:** Corresponding author: Koen van de Wetering, Department of Dermatology and Cutaneous Biology, Jefferson Institute of Molecular Medicine and PXE International Center of Excellence in Research and Clinical Care, Sidney Kimmel Medical College, Thomas Jefferson University, Philadelphia (PA), USA.

## Abstract

The membrane protein Ankylosis homologue (ANKH, mouse orthologue: ANK) prevents mineralization of joint-space and articular cartilage. The accepted view is that ANKH mediates cellular release of inorganic pyrophosphate (PPi), a strong physiological inhibitor of mineralization. Using global metabolite profiling, we identified citrate as the most prominent metabolite leaving HEK293 cells in an ANKH-dependent manner. Although PPi levels were increased in culture medium of HEK293-ANKH cells, PPi was formed extracellularly after release of ATP and other nucleoside triphosphates. *Ank*^*ank/ank*^ mice, which lack functional ANK, had substantially reduced concentrations of citrate in plasma and urine, while citrate was undetectable in urine of a human patient lacking functional ANKH. Bone hydroxyapatite of *Ank*^*ank/ank*^ mice also contained markedly reduced levels of citrate and PPi and displayed diminished strength. Together, our data show that ANKH is a crucial factor in extracellular citrate and PPi homeostasis that is essential for normal bone development.

## Introduction

Physiological mineralization is essential for normal development of vertebrates, but must be restricted to specific sites of the body. Vertebrates have evolved mechanisms to allow regulated mineralization in for instance bones and teeth, but prevent mineralization of soft connective tissues ^1,2^. The molecular details of the mechanism in vertebrates that restrict mineralization to specific sites of the body are incompletely characterized, however.

The *ANKH*/*Ank* (human/mouse) gene encodes a multi-span transmembrane protein involved in the prevention of pathological mineralization of cartilage and synovial fluid ^3,4^. *ANKH*/*Ank*, has a wide tissue distribution, which high levels of expression found in osteoblasts, prostate, skeletal muscle, brain and the cardiovascular system ^1,5,6^. A naturally occurring mouse mutant, progressive ankylosis (*Ank*^*ank/ank*^), presents early in life with progressive ankylosis of the spine and other joints, restricting mobility and critically limiting lifespan ^1^. Biallelic loss-of-function mutations in the human orthologue of *Ank*, *Ank homolog* (*ANKH),* underlie some forms of craniometaphyseal dysplasia (CMD), which also presents with progressive ankylosis, mainly affecting the spine and the joints of hands and feet ^7^. In 2000, Ho *et al.* showed that medium of *Ank*^*ank/ank*^ fibroblasts contained reduced concentrations of the physiological mineralization inhibitor inorganic pyrophosphate (PPi), leading to the now prevailing view that ANKH/ANK was the transport of PPi into the extracellular environment ^1,8^. An important source of extracellular PPi is ATP, which is extracellularly converted into AMP and PPi by membrane-bound ecto-nucleotidase pyrophosphatase/phosphodiesterase 1 (ENPP1) ^9^. We have previously shown that ATP release mediated by the hepatic membrane protein ATP-Binding Cassette subfamily C member 6 (ABCC6) is responsible for 60-70% of all PPi present in plasma ^10,11^.

Here we tested if release of ATP also underlies most of the PPi found in the extracellular milieu of ANKH-containing cells. Moreover, we applied global metabolite profiling ^12^ on medium of HEK293-ANKH cells to gain a comprehensive overview of metabolites extruded by cells in an ANKH-dependent manner. Our results provide new and unexpected insights into the substrate spectrum and anti-mineralization properties of ANKH and also show that ANKH has functions beyond inhibition of inhibition of pathological mineralization as it is, for instance, essential for normal bone development.

## Results

### HEK293-*ANKH* cells release ATP into the extracellular environment

To study the function of ANKH *in vitro*, we first generated several HEK293 cell lines overproducing wild type ANKH (ANKH^wt^) and ANKH^L244S^, a pathogenic loss-of-function mutant which still routes normally to the plasma membrane ^7^. As shown in Fig. 1A, endogenous ANKH was not detectable in parental HEK293 cells by immunoblot analysis, whereas high levels of ANKH protein were found in cells overexpressing *ANKH^wt^*. The loss-of-function *ANKH*^*L244S*^ mutant was also abundantly expressed, and a clone producing levels of the mutant protein higher than those detected in the HEK293-*ANKH*^*wt*^ cells was used for further analysis (Fig. 1A). First, we measured PPi levels in the medium of these cells over a 24-h time period and showed that PPi accumulated at higher levels in medium of HEK293-*ANKH^w^*^t^ cells than in medium of HEK293-*ANKH*^*L244S*^ or control HEK293 cells (Fig. 1B), confirming earlier reports that demonstrated the involvement of ANKH in extracellular PPi homeostasis ^1^. We have previously shown that ENPP1 produced by HEK293 cells converts extracellular ATP into AMP and PPi ^10^. Consequently, to determine what part of the PPi found in medium of *ANKH*^*wt*^ cells might be derived from extracellular ATP, converted by ENPP1 into AMP and PPi, AMP concentrations were quantified in the culture medium. As shown in Fig. 1C, a clear time-dependent increase in AMP concentrations was detected in medium of HEK293-*ANKH*^*wt*^ cells, while medium of untransfected HEK293 parental cells or cells producing the loss-of-function *ANKH*^*L244S*^ mutant contained only very little AMP. PPi and AMP concentrations in medium of *ANKH*^*wt*^ cells were within the same range (1-2 μM after 12 hours, compare panels B and C of Fig. 1) and the ratio of PPi to AMP was very similar to that previously reported for HEK293 cells overproducing ABCC6, a plasma membrane protein involved in the release of ATP ^10^. We attribute the somewhat lower abundance of AMP than PPi to further metabolism of AMP and the generation of PPi from other nucleoside triphosphates (NTPs) also released into the culture medium via ANKH (see below). A luciferase-based real-time ATP efflux assay was also carried out and confirmed that ANKH is involved in cellular ATP release (Figure 1D). Only HEK293-*ANKH*^*wt*^ cells showed robust ATP efflux, whereas release from HEK293-*ANKH*^*L244S*^ cells was indistinguishable from untransfected parental HEK293 cells in these assays. Collectively, these data indicate that HEK293-*ANKH*^*wt*^ cells release ATP, which is subsequently extracellularly converted into AMP and PPi.

**Fig 1.**
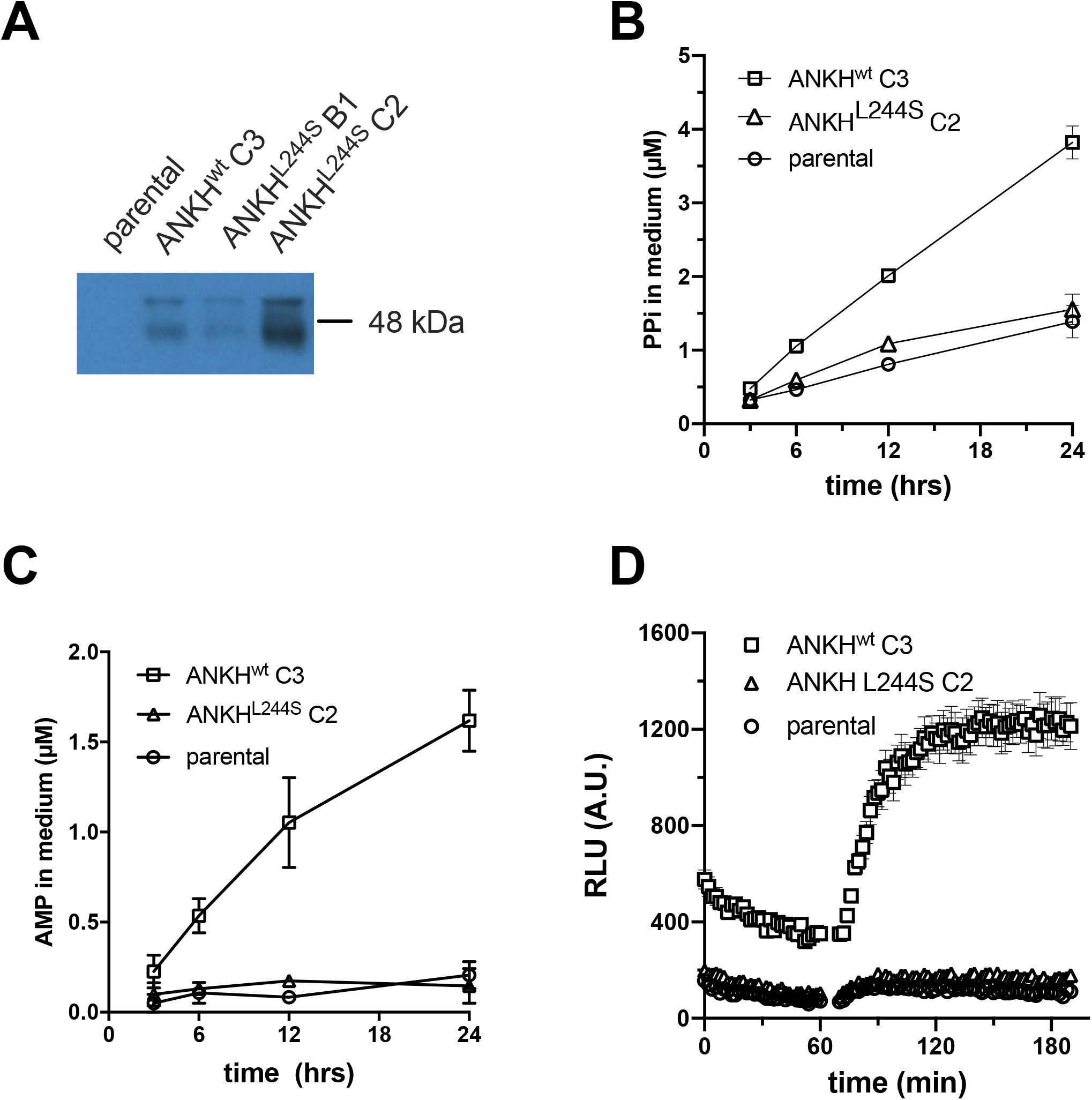
HEK293-*ANKH*^*wt*^ cells release ATP, which is rapidly converted into pyrophosphate (PPi) and AMP. Detection of ANKH in HEK293 parental, HEK293-*ANKH*^*wt*^ and HEK293-*ANKH*^*L244S*^ cells by immunoblot analysis (**A**). Concentrations of pyrophosphate (PPi) (**B**) and AMP (**C**) were quantified enzymatically and followed in medium samples of HEK293 parental, HEK293-ANKH^wt^ and HEK293-*ANKH*^*L244S*^ cells over the course of 24 hours. ATP release by HEK293 parental, HEK293-*ANKH*^*wt*^ and HEK293-*ANKH*^*L244S*^ cells was followed in real time using a luciferase-based assay (**D**). Results of representative experiments performed in triplicate are shown. In panels B and C data are expressed as mean +/− SD. Panel D shows mean +/− SEM.

### Culture medium of HEK293-*ANKH^wt^* cells contains large amounts of nucleoside monophosphates (NMPs)

In addition to ATP, ENPP1 can convert various other nucleoside triphosphates (NTPs) into their respective nucleoside monophosphate (NMP) and PPi. Our previous work has shown that ENPP1 activity in HEK293 cells is high ^10^. We therefore used liquid chromatography/mass spectrometry (LC/MS)-based global metabolite profiling to determine if ANKH also provides a pathway for release of other NTPs. Substantially elevated levels of AMP, CMP, GMP and UMP were detected in the culture medium of HEK293-*ANKH*^*wt*^ cells compared to untransfected parental and HEK293-*ANKH*^*L244S*^ cells (Fig. 2 A-D), For AMP and UMP differences between untranfected and HEK293-*ANKH*^*wt*^ cells reached statistical significance. These results support the hypothesis that ANKH provides a previously unanticipated pathway for cellular NTP release. Based on the levels of PPi, AMP and other NMPs detected in the culture medium, we estimate that cellular NTP release underlies at least 70% of the ANKH-dependent accumulation of PPi in the culture medium (for calculation see materials and methods section) of the PPi detected in medium of the HEK293-ANKH^wt^ cells.

**Fig 2.**
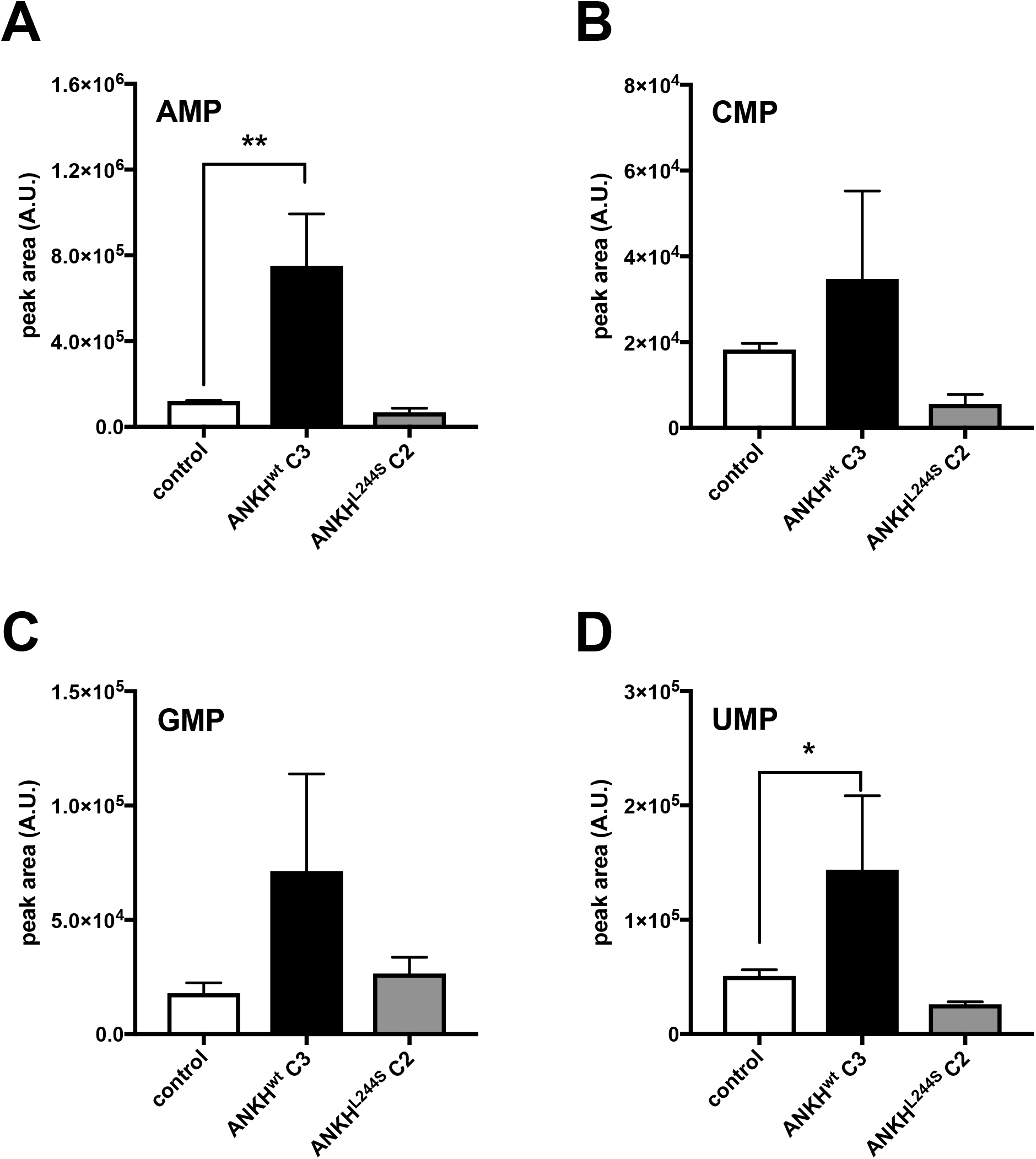
Medium of HEK293-*ANKH*^*wt*^ cells contains large amounts of nucleoside monophosphates. LC/MS-based global metabolite profiling was applied to 24-hour medium samples of HEK293 parental, HEK293-*ANKH*^*wt*^ and HEK293-*ANKH*^*L244S*^ cells. The relative abundance of masses corresponding to AMP (**A**), CMP (**B**), GMP (**C**) and UMP (**D**) were determined. Authentic standards were used to confirm the identity of NMPs. Data are expressed as mean +/− SD of an experiment performed in triplicate. * p < 0.05, ** p < 0.01.

### HEK293-ANKH cells release the TCA cycle intermediates citrate, succinate, and malate into the culture medium

The global metabolite profiling experiments also revealed that the calcium chelator citrate specifically accumulated in the culture medium of HEK293-*ANKH*^*wt*^ cells (Fig. 3A). Because global metabolite profiling experiments only provide relative metabolite levels, we also quantified citrate levels by LC/MS in 24-hour medium samples and found that approximately 1 mM citrate (2.5 μmol/24 hrs) was present in medium of HEK293-*ANKH*^*wt*^ cells, while it was almost undetectable in medium of HEK293 control and HEK293-*ANK*^*L244S*^ cells. To put this in perspective, the same medium samples of HEK293-*ANKH*^*wt*^ cells contained about 4 μM PPi (Fig. 1B), equivalent to the release of approximately 10 nmoles of NTPs. Thus, the amount of citrate released by the HEK293-*ANKH*^*wt*^ cells was at least 2 orders of magnitude higher than the amount of NTPs. Other metabolites found to be selectively elevated in medium of HEK293-*ANKH*^*wt*^ cells were malate (Fig. 3B) and succinate (Fig. 3C), although absolute levels increases relative to control cells were clearly less than those found for citrate. Using an independent enzymatic assay, citrate levels in culture medium were also followed over time and as shown in Fig. 3D, these experiments confirmed that citrate was present at approximately 1.1 mM in the 24-hour culture medium samples of the *ANKH*^*wt*^ cells, comparable to the concentration determined by LC/MS. Collectively these data show that ANKH is involved in the cellular release of large amounts of citrate.

**Fig 3.**
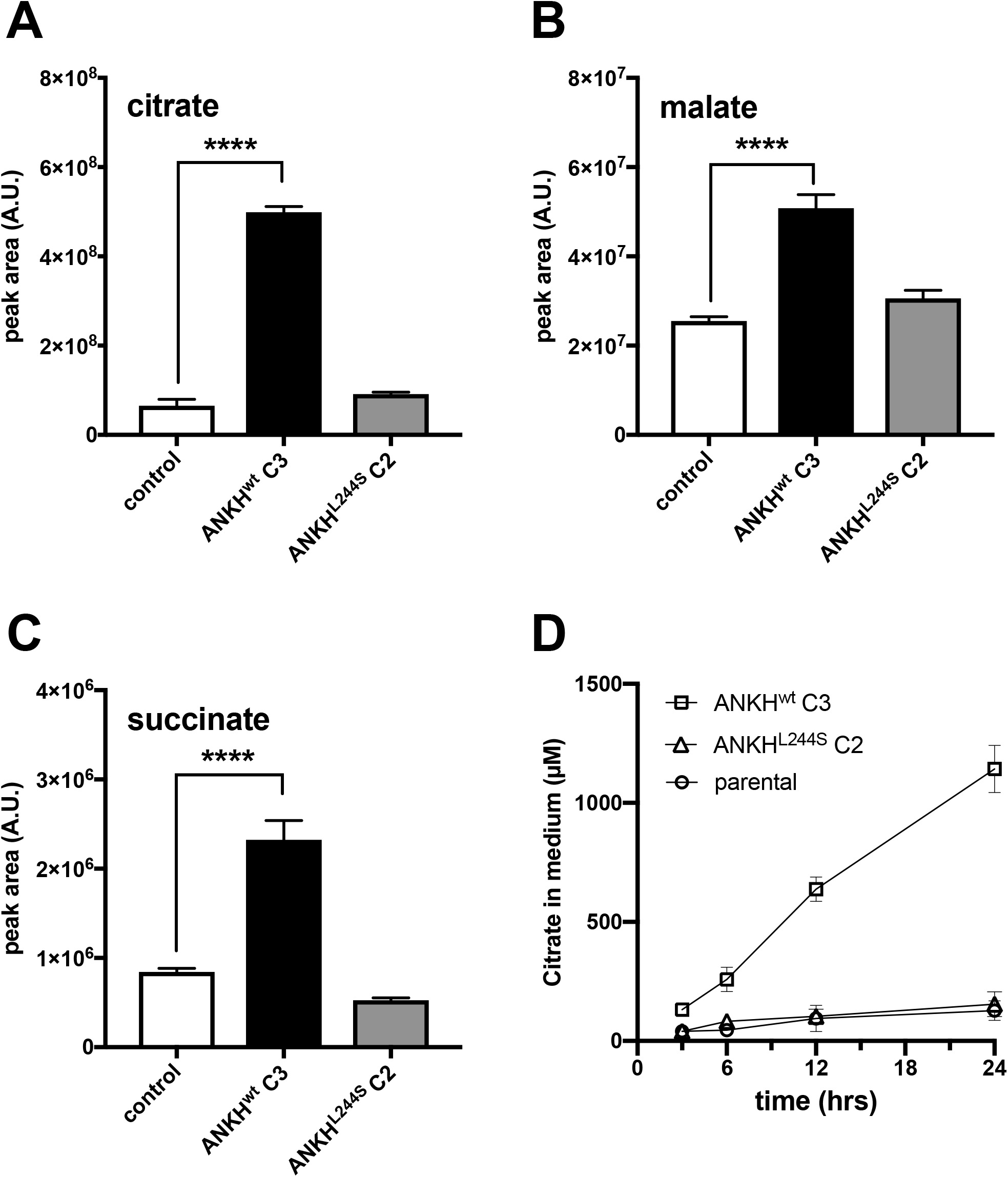
Medium of HEK293-*ANKH*^*wt*^ cells contains large amounts of citrate, succinate and malate. LC/MS-based global metabolite profiling was applied to 24-hour medium samples of HEK293 parental, HEK293-*ANKH*^*wt*^ and HEK293-*ANKH*^*L244S*^ cells. The relative abundance of masses corresponding to citrate (**A**), malate (**B**) and succinate (**C**) were determined. Authentic standards were used to confirm the identity of the Krebs-cycle intermediates. Using an enzymatic assay, citrate concentrations were followed for 24 hours in (**D**). Data are expressed as mean +/− SD of an experiment performed in triplicate. **** p < 0.0001.

### ANK affects PPi incorporation into bone

About 70% of the PPi found in plasma depends on ABCC6 activity ^11^, indicating that the contribution of ANKH/ANK to plasma PPi homeostasis is relatively minor. Consequently, instead of contributing to central PPi homeostasis in plasma, we hypothesized that ANKH/ANK is important in local PPi homeostasis. Osteoblasts express ANKH/ANK at relatively high levels ^5^ and the hydroxyapatite of bone contains substantial amounts of PPi ^13^. To determine if ANK has a role in incorporation of PPi in bone, we quantified PPi in tibiae and femora of wild type, *Ank*^*ank/ank*^, and mice heterozygous for *ank*. As shown in Fig. 4 PPi constituted about 0.1% (weight/weight) of bone tissue in wild type mice, whereas in *Ank*^*ank/ank*^ mice the amount of PPi associated with bone was reduced by approximately 75%. Moreover, in mice heterozygous for *ank*, PPi levels were also moderately (by approximately 25%), but significantly reduced. These data show that ANK is a crucial factor in PPi homeostasis in the local environment of bone tissue.

**Fig 4.**
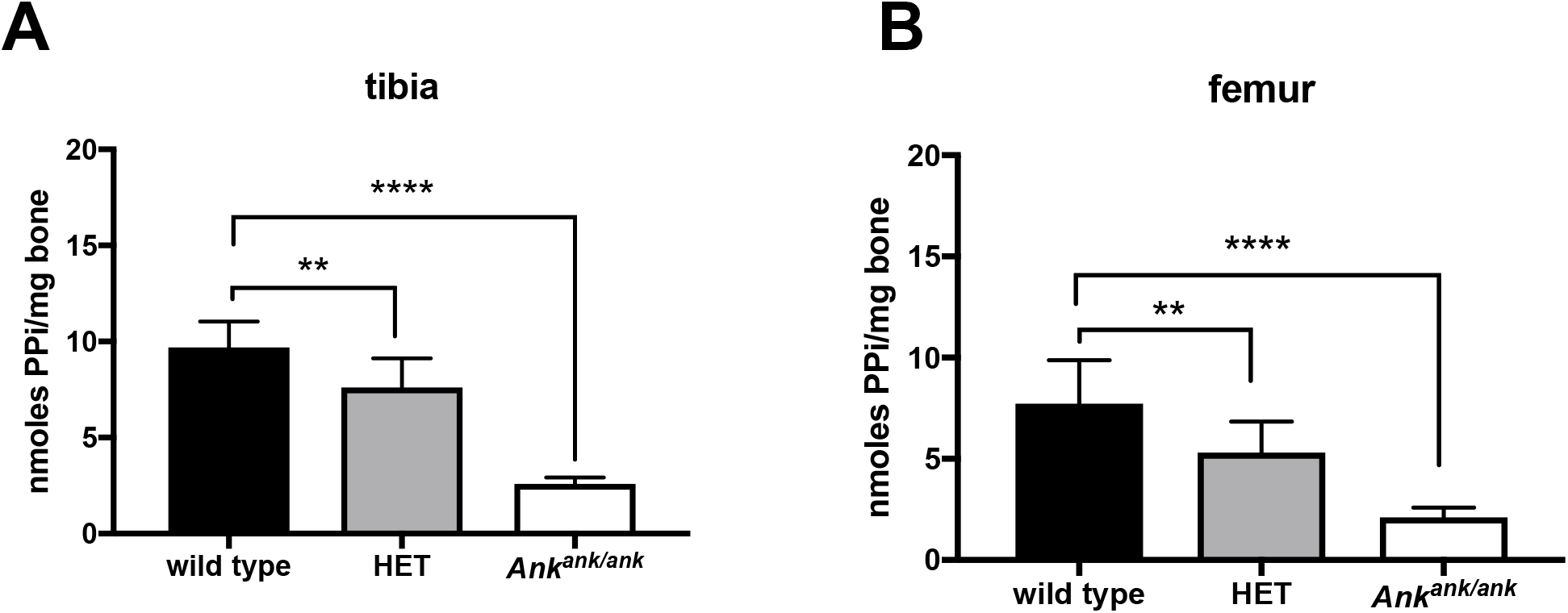
PPi content of bone tissue depends on ANK activity. Pyrophosphate content of tibiae (**A**) and femora (**B**) of wild type (n=10), heterozygous (HET, n=10) and *Ank*^*ank/ank*^ (*ank*/*ank*, n=8) mice. Data are expressed as mean +/− SD. ** p < 0.01, **** p < 0.0001.

### ANKH affects citrate disposition *in vivo*

Plasma contains substantial amounts of citrate ^14^. We therefore determined the effect of a complete inactivation of ANK in mice on plasma citrate concentrations and as shown in Fig. 5A, found that approximately 75% of citrate in plasma depended on ANK. Because citrate is also one of the most abundant organic anions in urine ^15^, we measure citrate excretion in *Ank*^*ank/ank*^ mice. As shown in Fig. 5B, the *ank* mutant mice excreted approximately 40% less citrate via their urine than their wild type litter mates. The availability of an NMR spectrum of urine of a 19-year-old female CMD patient carrying biallelic homozygous inactivating mutations in *ANKH* (*ANKH^L244S^*), previously described by Morava *et al.* ^7^ made it possible to carry out an analysis of citrate levels. Citric acid was not detected in urine of this CMD patient (Fig. 5C, upper panel). The lower panel of Fig. 5C shows the typical citrate resonance in urine of a representative age-matched control, which contained 370 μmol citrate/mmol creatinine. It is interesting to note that the succinate resonance is visible in the NMR spectrum of control urine, while its concentration is clearly much lower in urine of the CMD patient (Fig. 5E). These data suggest that ANKH impacts the *in vivo* disposition of succinate and especially citrate in both, humans and mice.

**Fig 5.**
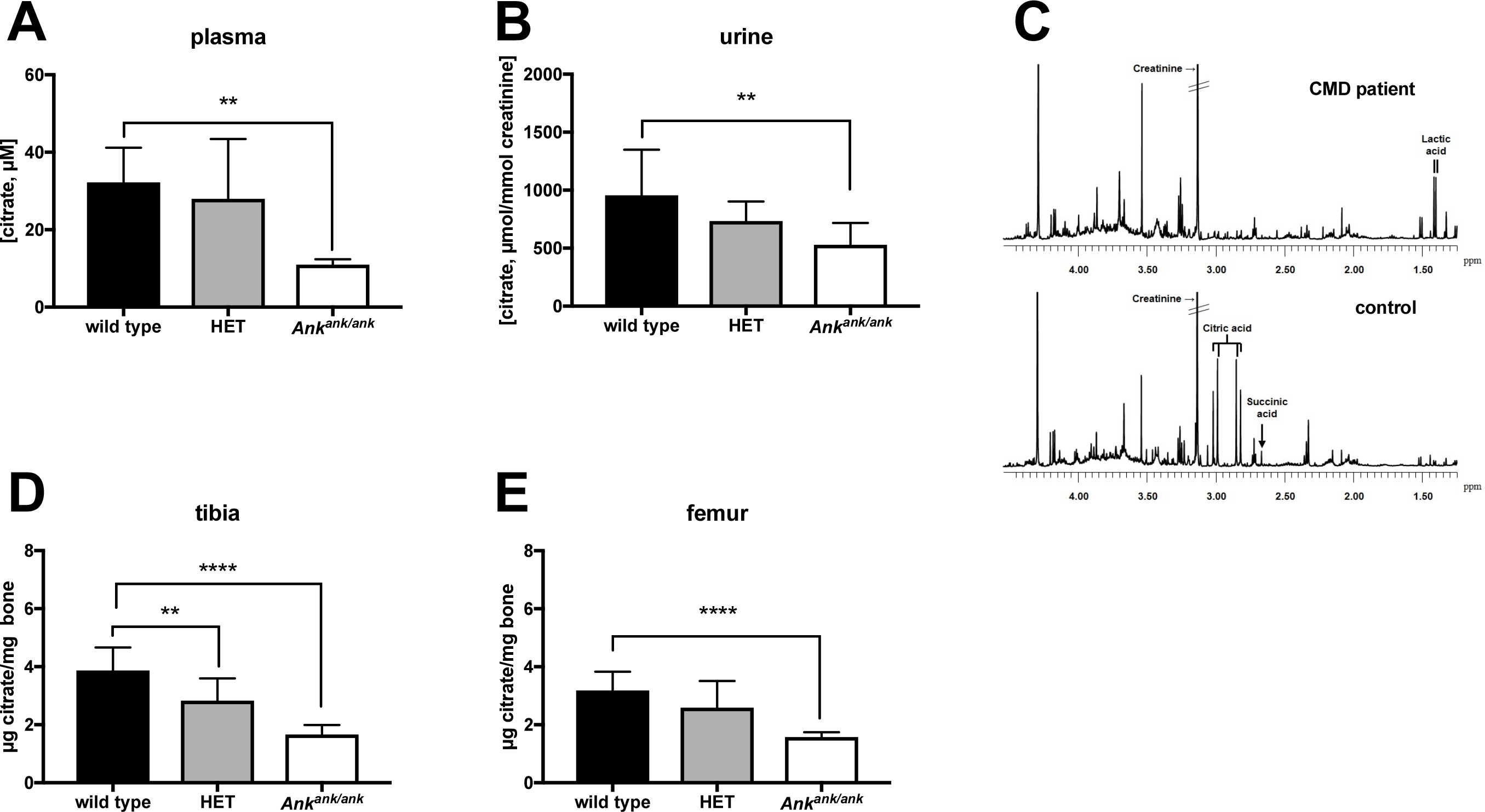
Extracellular Citrate depends on ANK/ANKH activity. (**A**) Citrate plasma concentrations in wild type (n=8), heterozygous (HET, n=8) and *Ank*^*ank/ank*^ (n=8) mice. Citrate concentrations in urine of wild type (n=6), heterozygous (HET, n=10) and *Ank*^*ank/ank*^ (n=9) mice. (**C)**Urine of a patient suffering from craniometaphyseal dysplasia (CMD) due to biallelic inactivating mutations in *ANKH* is virtually devoid of citrate. NMR spectra of urine of a patient with biallelic pathogenic *ANKH*^L244S^ mutations (C, upper panel). A representative sex- and age-matched control urine sample contained 370 μmol/mmol creatinine (**C,** lower panel). Spectra are scaled on creatinine. Citrate resonates as a typical AB-system (2.98 ppm; four peaks between 2.80 and 3.05 ppm). Reference values for urinary citrate for this age group are 208-468 μmol/mmol creatinine (n=20 healthy controls) ^36^. Succinate resonates as a singlet resonance at 2.66 ppm. For unknown reasons, urinary lactate was somewhat increased in urine of the CMD patient (120 μmol/mmol creatinine; reference <75 μmol/mmol creatinine). Citrate content of tibiae (**D**) and femora (**E**) of wild type (n=10), heterozygous (HET, n=10) and *Ank*^*ank/ank*^ (n=10) mice. Data are expressed as mean +/− SD. * p < 0.05, ** p < 0.01, *** p< 0.001, **** p < 0.0001.

Like PPi, citrate is also one of the major organic compounds present in bone and also strongly associates with hydroxyapatite ^16^. With 90% of the body’s citrate content present in bone, this tissue is thought to play a central role in extracellular citrate homeostasis ^17^. Therefore, we determined if bone citrate levels depend on ANK. These experiments revealed that femora and tibiae of *Ank*^*ank/ank*^ mice contained approximately 50% less citrate than the same bones of wild type mice (Fig. 5D,E). Moreover, bones of mice heterozygous for *ank* also contained less citrate, which in the case of tibia was significantly lower than in wild type mice (Fig. 5D). Together these data attest to the major impact of ANK on citrate homeostasis of bone.

### Material properties of bone tissue of *Ank*^*ank/ank*^ mice are altered

We next explored the consequences of the absence of ANK activity on bone physiology, by characterizing geometry and density of femurs harvested from *Ank*^*ank/ank*^, wild type and mice heterozygous for *ank* by microCT. At 3 months of age, most of the bone parameters, including bone area (Fig. 6A), tissue mineral density (TMD, Fig. 6B), and cortical thickness (Fig. 6C), were not significantly different between wild type and *Ank*^*ank/ank*^ mice. However, significant differences in cortical bone properties between *Ank*^*ank/ank*^ and wild type mice were detected for bone area fraction (−12.1%), cortical bone perimeter (+9.8%), and cross-sectional geometry as indexed by eccentricity (−9.4%). Next, the structural and material properties of the bone were determined by standard three-point bending. Plotting ultimate bending moment against section modulus (Fig. 6G) yielded linear relationships for each genotype (r^2^ = 0.84 wild type, 0.73 HET, 0.67 *Ank*^*ank/ank*^) that did not significantly differ in slope (p = 0.88). However, we observed that femurs from *Ank*^*ank/ank*^ mice required significantly less force per equivalent area of bone to break, as demonstrated by a significant difference in regression intercept (p = 0.0170). Taken together, our results indicate that the geometry of femora of *Ank*^*ank/ank*^ mice is altered and that these femora have diminished whole bone strength per equivalent amount of bone, results that are consistent with published data showing citrate deposition in bone affects hydroxyapatite nanostructure and strength ^16^.

**Fig 6.**
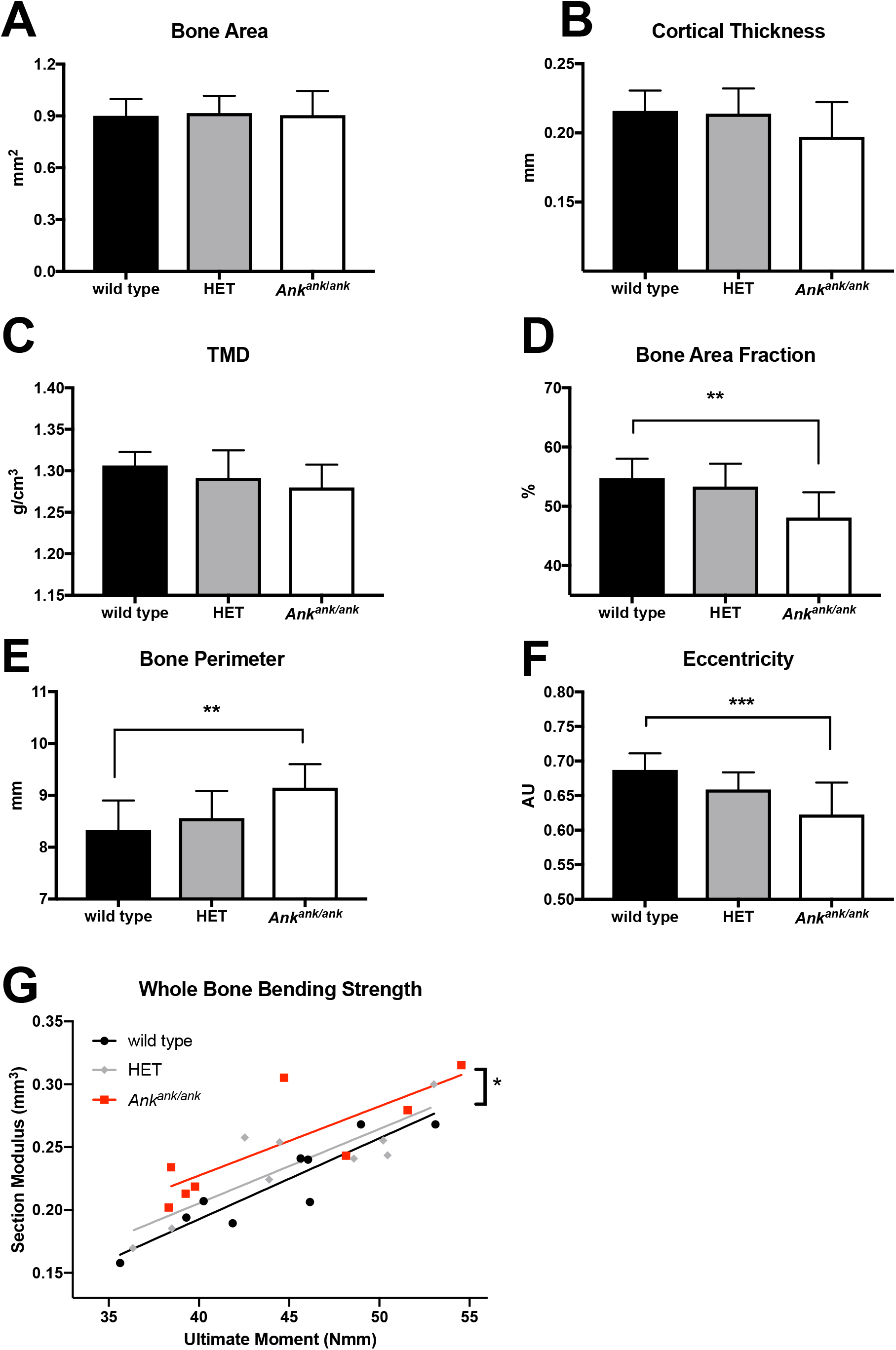
Bone geometry and mechanical performance is altered in the absence of ANK activity. microCT was used to determine (A) bone area, (B) cortical thickness, (C) tissue mineral density (TMD), (D) bone area fraction (B.Ar/T.Ar), (E) bone perimeter, and (F) eccentricity in femora of wild type (n=9), heterozygous (HET, n=10) and *Ank*^*ank/ank*^ (n=8) mice. (G) To compare whole bone bending strength, a linear regression between section modulus and ultimate bending moment was analyzed for each genotype (r^2^ = 0.84 wild type, 0.73 HET, 0.67 *Ank*^*ank/ank*^). The slope was not different between genotypes, but the intercept was significantly different in femora from *Ank*^*ank/ank*^ mice, which utilized an increased section modulus to achieve the corresponding ultimate moment. * p < 0.05, ** p < 0.01, *** p < 0.001.

## Discussion

Ectopic mineralization – the deposition of hydroxyapatite in soft connective tissues – can be a sequela of a number of clinical conditions, including aging, cancer, diabetes, chronic kidney disease and genetic disorders. With no effective treatment currently available, ectopic mineralization is associated with significant morbidity and mortality ^18^.

ANKH/ANK is known for its important role in the prevention of pathological mineralization of joints, and its absence results in severe, progressive, ankylosis in both, humans and mice. It was previously thought that the main function of ANKH/ANK lies in regulation of extracellular PPi homeostasis, but here we identified a new and previously unanticipated function of ANKH/ANK: regulation of extracellular citrate concentrations. Although citrate has long been known to be a major compound in plasma, urine and bone, the mechanism used by cells to extrude citrate has been elusive. Our current data firmly link a specific protein, ANKH/ANK to extracellular citrate disposition *in vivo*. Notably, our results are in line with previous GWAS studies describing a correlation between plasma citrate levels and certain *ANKH* variants in humans ^19^ and an association found in cows between intronic *ANKH* variants and milk citrate concentrations ^20^.

Extracellular citrate is present in many tissues and body fluids where is serves diverse and, in some cases, unknown functions ^14^. In human plasma citrate levels are substantial (100-300 μM), but its function is unclear ^21^. Various cell types express citrate uptake transporters and it has been proposed that plasma citrate provides an additional energy source for cells under hypoglycemic conditions ^14^. Alternatively, citrate is a powerful anticoagulant and might prevent pathological blood clotting.

Via glomerular filtration, plasma citrate ends up in urine, where it contributes to the prevention kidney stone formation ^22^. Whereas urine of the *Ank*^*ank/ank*^ mice still contained substantial amounts of citrate, that of the human CMD patient lacking functional ANKH was virtually devoid of citrate. This difference might be partly explained by dietary differences: Citrate has a high bioavailability of 80-90% ^23^ and is present in standard rodent food. Possibly, the human CMD patient had a diet that was low in citrate, whereas part of the citrate detected in plasma of *Ank*^*ank/ank*^ mice comes from dietary sources.

Most of the body’s citrate, over 90%, is present in bone tissue, where it stabilizes hydroxyapatite^16^. Our results show that about 50% of bone citrate depends on ANK activity, in line with the high expression of *Ank* in osteoblasts ^5^. The altered material properties of *Ank*^*ank/ank*^ bones, *i*.*e*. the altered relationship between ultimate moment and section modulus, nicely fits the described role of citrate in stabilizing hydroxyapatite. The altered eccentricity and perimeter of *Ank*^*ank/ank*^ femora are most likely a result of compensatory bone remodeling to retain whole bone strength without increasing total bone mass. Interestingly, Ma *et al.* recently reported that local levels of extracellular citrate are also important for the osteogenic development of human mesenchymal stem cells ^17^. It is therefore conceivable that ANKH-dependent citrate release into bone is not only important for the material properties of hydroxyapatite, but also contributes to osteogenic differentiation.

Relatively high extracellular citrate concentrations of approximately 400 μM are found in the brain. Astrocytes actively release citrate, which is used by neurons as energy source under hypoglycemic conditions ^14^. Patients suffering from CMD due to inactivating mutations in *ANKH* suffer from mental retardation ^7^, suggesting a function of ANKH in brain physiology. The highest extracellular citrate concentrations are found in prostatic fluid (up to 180 mM). ANKH is expressed at high levels in the epithelial cells of the prostate, known to release citrate into prostatic fluid. Although a specific splice variant of the mitochondrial citrate carrier SLC25A1 has been implied in citrate efflux from the prostate ^24^, ANKH likely contributes to this process. In summary, extracellular citrate is present in many tissues and body fluids and we anticipate that our discovery that ANKH/ANK is involved in extracellular citrate homeostasis will allow clarifying its function in other tissues for instance by using the Ank^*ank/ank*^ mouse model.

A second important finding of the current study is that most, if not all, PPi found in the extracellular environment of ANKH/ANK containing cells, originates from released NTPs, which are extracellularly converted into their respective NMP and PPi by ENPP1. This contradicts earlier work, proposing direct ANKH/ANK-dependent cellular efflux of PPi ^1^. Our current data strongly support the conclusion that ANKH/ANK mediates release of NTPs release, not PPi. First, *in vitro* experiments showed that the majority of PPi found in the culture medium of HEK293-*ANKH*^*wt*^ cells was derived from NTP efflux. Two reports have appeared that suggest cells release ATP in an ANKH-dependent manner ^25–27^. These studies did however not quantify the relative amounts of extracellular ATP, AMP and PPi and therefore did not allow assessment of the relative contribution of ANKH-mediated ATP release to extracellular PPi concentrations. Strong evidence arguing against direct PPi transport by ANKH/ANK also comes from our analysis of bones of mice lacking ENPP1. Moreover, 75% of the PPi present in bone depends on ANK activity (Fig. 4). If ANK would directly transport PPi, incorporation of this fraction into bone would not require ENPP1 activity. However, we found that PPi is virtually absent in bones of *Enpp1*^−/−^ mice (*asj^GrsrJ^*) (Szeri et al, manuscript in preparation), which is only compatible with ANKH/ANK mediating NTP release with subsequent extracellular formation of PPi by ENPP1. The function of PPi in bone tissue is not completely clear, but might be related to stabilization of hydroxyapatite and, consequently, bone mineral density. Such a function would fit data of previous studies showing that bones of *Enpp1*^−/−^ mice, which virtually lack PPi (Szeri et al, manuscript in preparation), have a substantially greater reduction in mineral density ^28,29^ than bones of *Ank*^*ank/ank*^ mice. These data also indicate that the residual 25% of PPi found in bones of *Ank*^*ank/ank*^ mice suffices to a large extent to keep BMD close to the normal range. The proposed effects of PPi on mineral density are similar to the effects of bisphosphonates, pharmaceutical PPi analogues that are widely used in the treatment of osteoporosis ^30^. Kim et al ^5^ have previously found a more dramatic effect of ANK on bone mineral density. A different genetic background of their *Ank*^*ank/ank*^ mice might underlie this more dramatic effect. Food composition, specifically varying PPi content ^31^, might also have contributed to the differences found in BMD between the two studies.

ANKH/ANK has previously been shown to inhibit ectopic mineralization in the microenvironment of the joint space ^1^. Low plasma levels of PPi underlie several other genetic mineralization disorders ^18,32^. Plasma citrate levels depend on ANK activity, demonstrating that ANK substrates end up in the blood circulation. This does not come as a surprise given the wide tissue distribution of ANKH/ANK ^1^. Most likely, NTPs are also released into the blood circulation via ANK, allowing subsequent PPi formation in plasma. Under normal conditions, 60-70% of plasma PPi comes from ABCC6-mediated hepatic NTP secretion ^10,11^. ANK can therefore be expected to be responsible for part of the remaining 30-40% of the PPi present in plasma that is independent of ABCC6 activity. The relatively small contribution of ANK together with the large variability in plasma PPi concentrations ^10,11,31^ prevents determination of the contribution of ANK to plasma PPi in *Ank*^*ank/ank*^ mice. *Ank^ank/ank^*;*Abcc6*^−/−^ compound mutant mice (*Ank^ank^*;*Abcc6^tm1Jfk^*) provide the optimal experimental model system to determine the contribution of ANKH/ANK to plasma PPi. If ANKH indeed contributes to plasma PPi homeostasis, it represents an attractive pharmacological target in ectopic mineralization disorders caused by low plasma levels of PPi. For instance, stimulation of ANKH activity in patients suffering from pseudoxanthoma elasticum (PXE), a slowly progressive ectopic calcification disorder caused by inactivating mutations in the gene encoding the hepatic efflux transporter ABCC6 ^32^, might increase plasma PPi concentrations and halt disease progression. As citrate chelates calcium and has been shown to prevent kidney stone (uroliths) formation ^22^, ANKH-mediated citrate release might also contribute to inhibition of ectopic mineralization in joints and other tissues. The previous observation of *Ho et al*. ^1^ that *Ank*^*ank/ank*^ mice have an increased incidence of kidney calcification would fit a function of ANKH/ANK in prevention of ectopic mineralization in tissues different from those lining the joints.

In conclusion, we identified ANKH/ANK as an important player in cellular release of citrate and NTPs. Citrate might have a previously unanticipated role in the prevention of soft tissue mineralization, in addition to other major ectopic mineralization inhibitors like PPi, Mg^2+^ and Fetuin-A ^33,34^. Moreover, we found that ANKH/ANK is a crucial factor in normal bone physiology by determining the amount of citrate and PPi incorporated in bone tissue.

## Materials and methods

### Cell culture

HEK293 cells were passaged in HyClone DMEM (GE) supplemented with 5% FBS and 100 units pen/strep per ml (Gibco) at 37°C and 5% CO_2_ under humidified conditions. Efflux experiments were performed in 6-well plates in 2.5 ml Pro293a medium (Lonza) supplemented with 2 mM L-glutamine and 100 units pen/strep per ml.

### Animals

Mice heterozygous for the progressive ankylosis allele (*ank*) were obtained from The Jackson Laboratory (Bar Harbor, ME; C3FeB6 *A/A^w-J^-Ank^ank/J^*, stock number 000200). Heterozygote breeders were used to generate *Ank*^*ank/ank*^, heterozygous and wild type littermates. Protocols were approved by the Institutional Animal Care and Use Committee of Thomas Jefferson University in accordance with the National Institutes of Health Guide for Care and Use of Laboratory Animals. Animals analyzed were between 11-14 weeks old. Plasma samples were collected by cardiac puncture in heparinized syringes. Studies included similar numbers of male and female mice.

### Mutagenesis and overexpression of ANKH

*ANKH*^*wt*^ cDNA was obtained from Sino Biological and subcloned into pEntr223 by USER cloning. The *L244S* mutation was introduced by USER cloning with primers 5’-ACCAGAAGC<u>U</u>CAGCATCTTTCTTATTGTTGCATCTCCC-3’ and AGCTTCTGG<u>U</u>GGCCTTCCGCTC TAATTCTGGCCACA. cDNAs were subsequently subcloned in a Gateway compatible pQCXIP expression vector ^10^. HEK293 cells were transfected with pQCXIP-ANKH by calcium phosphate precipitation. ANKH^wt^ and ANKH^L244S^ in clones resistant to 2 μM puromycin were determined by immunoblot analysis, with a polyclonal antibody directed against ANKH (OAAB06341, Aviva Systems Biology).

### Enzymatic quantification of PP_i_, AMP and citrate

In medium samples, PP_i_ and AMP were quantified as described ^11^ with modifications. PPi concentrations were determined using ATP sulfurylase from NEB, and adenosine 5’phosphosulfate from Cayman Chemicals. AMP was quantified as follows: To 1 μl of sample or standard, 100 μl of a solution containing 0.14 U/ml pyruvate orthophosphate dikinase (PPDK, kind gift of Kikkoman Chemifa), 12.5 μmol/L PPi (Sigma-Aldrich), 40 μmol/L phosphoenol pyruvate (Cayman Chemicals), 50 μmol/L dithiothreitol, 1 mmol/L EDTA, 7.5 mmol/L MgSO_4_ and 30 mmol/L BES (pH 8.0) was added. Conversion of AMP into ATP was allowed to proceed for 20 min at 45 °C, after which PPDK was inactivated by incubation at 80 °C for 10 min. To determine PPi and citrate amounts in bones, tibiae and femora of 13-week-old mice were collected and defleshed. Epiphyses were removed and bone marrow was spun out of the bones (30,000 RCF, 1 min). Bones were subsequently dissolved by incubation with continuous mixing in 10% formic acid (60 °C, 750 RPM, 14 hrs). Samples were spun for 10 min at 30,000 RCF and the supernatant was analyzed for PPi and citrate content. For bone extracts a slightly modified, more sensitive, version of the PPi assay was used. A total reaction volume of 520 μl assay mix contained 100 μl of SL-ATP detection reagent (Biothema, Sweden), 0.1 μl ATP removal reagent (“apyrase”, BioThema, Sweden), 6 μM adenosine-5’-phosphosulphate (APS) (SantaCruz, TX), 0.15 U/ml ATP sulphurylase (ATPS) (New England Biolabs) and 400 μl of ATP-free Tris-EDTA buffer (BioThema, Sweden) was first incubated overnight at room temperature to convert PPi into ATP for subsequent degradation by apyrase. The overnight incubation removed background PPi from the assay mixture, resulting in a higher sensitivity of the assay. Next, the sample, diluted 500-fold in Tris-EDTA buffer, was added to 500 μl of the assay mixture, resulting in an increase in luminescence due to the conversion of PPi and APS into ATP, a reaction catalyzed by ATPS. Finally, a known amount of ATP was added as internal standard and the ratio between the increase in bioluminescent signal induced by the addition of PPi and by the increase induced by the addition of ATP was used to calculate the PPi concentration. The assay was performed in a Berthold FB12 luminometer in the linear range of the detector. Internal PPi standards were used to show robustness and sensitivity of the assay.

Citrate was quantified in medium samples using the Megazyme Citric Acid Kit (Megazyme, Ireland).

### Real-time ATP efflux assay

Real-time ATP efflux assays were performed as described ^11^, with modifications. To reduce ATP release by the initial buffer change, cells were incubated at 27°C, for 1 hr. Then an additional 50 μl of ATP efflux buffer containing 10% of ATP-monitoring reagent (BactiterGlo, Promega), dissolved in ATP efflux buffer was added. Bioluminescence was followed in real-time for 1 hr at 27 °C and 2 hrs at 37 °C in a Flex Station3 microplate reader (Molecular Devices).

### LC/MS-based global metabolite profiling

Proteins were precipitated in 200 μl of medium or 50 μl plasma by adding 800 μl and 200 μl acetonitrile:methanol (1:1), respectively. Samples were shaken (10 minutes, 500 RPM, 21°C), centrifuged (15,000 g, 4°C, 10 min) and the supernatant dried in a Speed-Vac. Pellets were stored at −20°C until analysis. For analysis pellets were suspended in 45 μl mobile phase A of which 10 μl was analyzed by ion-pairing LC/MS as described ^12^.

Analytes were identified based on accurate mass and retention time, which matched reference standards. Peak areas were determined using Masshunter Qualitative Analysis software version 7.0SP2 (Agilent Technologies).

### LC/MS-based quantification of citrate

Plasma proteins were removed as described above and resuspended in 50 ul mobile phase A, while urine and bone samples were diluted in mobile phase A (5 and 20-fold, respectively). A volume of 5 μl of each sample was analyzed as described under LC/MS global metabolite profiling, along with calibration curves consisting of mobile phase A spiked with citrate concentrations ranging from 1 to 1000 μM. Quantification was performed using Masshunter Profinder Quantitative Analysis software version B.08.00, service pack 3 (Agilent Technologies).

### NMR spectroscopy

One-dimensional ^1^H-NMR spectroscopy of urine samples was performed as described ^35^. Briefly, urine samples were centrifuged for 10 min at 3,000 g and trimethylsilyl-2,2,3,3-tetradeuteropropionic acid (TSP; sodium salt; Sigma) in D_2_O was added before analysis to serve both, as an internal quantity reference and a chemical shift reference. The pH of each sample was adjusted to 2.50 ± 0.05 with concentrated HCl. ^1^H-NMR spectra were obtained using a Bruker 500-MHz spectrometer (pulse angle: 90°; delay time: 4 s; no. of scans: 256; relaxation delay: 2s). Assignment of peak positions for compound identification was performed by comparing the peak positions in the spectra of the metabolites with the reference spectral database of model compounds at pH 2.5 using Amix version 3.9.14 (Bruker BioSpin).

### Calculation of the contribution of NTP release to ANKH-dependent accumulation of PPi in the culture medium

To estimate the contribution of ANKH^wt^-mediated NTP release to 24-hour extracellular PPi concentrations, PPi concentrations in medium of HEK293 parental cells were subtracted from the PPi concentrations detected in medium of HEK293-*ANKH*^*wt*^ cells, yielding an ANKH-specific PPi accumulation in the 24-hr culture medium samples of 2.4 μM. The same calculation demonstrated an ANKH-specific accumulation of 1.4 μM AMP in the culture medium. This demonstrated that ATP release underlies at least 60% of the ANKH-dependent PPi accumulation detected in the culture medium (1.4/2.4 × 100 = 58). GMP, UMP and CMP were also found to increase in culture medium in an ANKH-dependent manner. Based on the relative LC/MS signals of the NMPs, we estimated that AMP was responsible for 80% of the total NMP concentration in the culture medium, whereas GMP, UMP and CMP together were responsible for the remaining 20%. Together these data demonstrate that nucleoside monophosphate (NMP) concentrations could explain 70% of the ANKH-dependent PPi that had accumulated in the culture medium after 24 hrs. The calculated 70% is most likely an underestimation, as generated NMPs will be further metabolized by the HEK293 cells, as we have observed before ^10^.

### MicroCT

Each bone was scanned using a Bruker Skyscan 1275 microCT system equipped with a 1 mm aluminum filter. One femur from each mouse was scanned at 55 kV and 181 μA with a 74 ms exposure time. Transverse scan slices were obtained by placing the long axis of the bone parallel to the z axis of the scanner using a custom 3D printed sample holder. An isometric voxel size of 13 μm was used. Images were reconstructed using nRecon (Bruker) and analyzed using CTan (Bruker).

### Three-point Bending assay

Three-point bending was performed on bones that had been stored at −20 °C in PBS-soaked gauze after harvest. Femora were scanned with microCT before performing three-point bending. Briefly, each femur was oriented on a standard fixture with femoral condyles facing down and a bending span of 8.7 mm. Next, a monotonic displacement ramp of 0.1 mm/s was applied until failure, with force and displacement acquired digitally. The force-displacement curves were converted to stress-strain using microCT-based geometry and analyzed using a custom GNU Octave script.

### Statistical analyses

P-values of group comparisons were calculated using one-way Anova using Prism 7.0d version (GraphPad Software Inc.). Significance is indicated in the figures, with * < 0.05, ** < 0.01, *** < 0.001 and **** < 0.0001.

## Author contributions

FS Wrote manuscript, performed experiments

SL Performed experiments

SD Performed experiments

UE Performed NMR analysis of urine samples

RE Provided access to essential equipment

KR Provided access to critical equipment

CW Provided essential reagents

JPS Provided essential reagents

RW Provided patient sample and analyzed data

RET performed experiments, analyzed data

RJ Performed metabolomics analyses, analyzed data

KvdW Wrote manuscript, analyzed data, conceptualized project, supervised project

All Authors have seen and reviewed the manuscript

## Acknowledgements

We thank our colleagues Piet Borst (The Netherlands Cancer Institute), Susan Cole (Queens’s University) and Jouni Uitto (Thomas Jefferson University) for critically reviewing our manuscript and valuable discussions. FSz received financial support from the Fulbright Visiting Scholar Program sponsored by the U.S. Department of State and a mobility grant from the Hungarian Academy of Sciences. Further funding for this work was provided by PXE International and the National Institutes of Health Grant R01AR072695 (KvdW).

## Competing Interests

The authors declare they have no financial or non-financial competing interests.

## References

1. Ho, A. M., Johnson, M. D. & Kingsley, D. M. Role of the mouse ank gene in control of tissue calcification and arthritis. Science 289, 265–270 (2000).

2. Kawasaki, K., Buchanan, A. V. & Weiss, K. M. Biomineralization in humans: making the hard choices in life. Annu. Rev. Genet. 43, 119–142 (2009).

3. Gurley, K. A. et al. Mineral formation in joints caused by complete or joint-specific loss of ANK function. J. Bone Miner. Res. 21, 1238–1247 (2006).

4. Abhishek, A. & Doherty, M. Pathophysiology of articular chondrocalcinosis--role of ANKH. Nat. Rev. Rheumatol. 7, 96–104 (2011).

5. Kim, H. J., Minashima, T., McCarthy, E. F., Winkles, J. A. & Kirsch, T. Progressive ankylosis protein (ANK) in osteoblasts and osteoclasts controls bone formation and bone remodeling. J. Bone Miner. Res. 25, 1771–1783 (2010).

6. Wu, C. et al. BioGPS: an extensible and customizable portal for querying and organizing gene annotation resources. Genome Biol. 10, R130–8 (2009).

7. Morava, E. et al. Autosomal recessive mental retardation, deafness, ankylosis, and mild hypophosphatemia associated with a novel ANKH mutation in a consanguineous family. J. Clin. Endocrinol. Metab. 96, E189–98 (2011).

8. Orriss, I. R., Arnett, T. R. & Russell, R. G. G. Pyrophosphate: a key inhibitor of mineralisation. Curr. Opin. Pharmacol. 28, 57–68 (2016).

9. Nitschke, Y. & Rutsch, F. Inherited Arterial Calcification Syndromes: Etiologies and Treatment Concepts. Curr. Osteoporos. Rep. 15, 1–16 (2017).

10. Jansen, R. S. et al. ABCC6 prevents ectopic mineralization seen in pseudoxanthoma elasticum by inducing cellular nucleotide release. Proc. Natl. Acad. Sci. U.S.A. 110, 20206–20211 (2013).

11. Jansen, R. S. et al. ABCC6-mediated ATP secretion by the liver is the main source of the mineralization inhibitor inorganic pyrophosphate in the systemic circulation-brief report. Arterioscler. Thromb. Vasc. Biol. 34, 1985–1989 (2014).

12. Goncalves, M. D. et al. High-fructose corn syrup enhances intestinal tumor growth in mice. Science 363, 1345–1349 (2019).

13. Alfrey, A. C. & Solomons, C. C. Bone pyrophosphate in uremia and its association with extraosseous calcification. J. Clin. Invest. 57, 700–705 (1976).

14. Mycielska, M. E., Milenkovic, V. M., Wetzel, C. H., Rümmele, P. & Geissler, E. K. Extracellular Citrate in Health and Disease. Curr. Mol. Med. 15, 884–891 (2015).

15. Petrarulo, M., Facchini, P., Cerelli, E., Marangella, M. & Linari, F. Citrate in urine determined with a new citrate lyase method. Clin. Chem. 41, 1518–1521 (1995).

16. Hu, Y.-Y., Rawal, A. & Schmidt-Rohr, K. Strongly bound citrate stabilizes the apatite nanocrystals in bone. Proc. Natl. Acad. Sci. U.S.A. 107, 22425–22429 (2010).

17. Ma, C. et al. Citrate-based materials fuel human stem cells by metabonegenic regulation. Proc. Natl. Acad. Sci. U.S.A. 115, E11741–E11750 (2018).

18. Uitto, J., Li, Q., van de Wetering, K., Váradi, A. & Terry, S. F. Insights into Pathomechanisms and Treatment Development in Heritable Ectopic Mineralization Disorders: Summary of the PXE International Biennial Research Symposium-2016. J. Invest. Dermatol. 137, 790–795 (2017).

19. Shin, S.-Y et al. An atlas of genetic influences on human blood metabolites. Nat. Genet. 46, 543–550 (2014).

20. Sanchez, M.-P. et al. Sequence-based GWAS, network and pathway analyses reveal genes co-associated with milk cheese-making properties and milk composition in Montbéliarde cows. Genet. Sel. Evol. 51, 34–19 (2019).

21. Rudman, D. et al. Hypocitraturia in patients with gastrointestinal malabsorption. N. Engl. J. Med. 303, 657–661 (1980).

22. Raffin, E. P. et al. The Effect of Thiazide and Potassium Citrate Use on the Health Related Quality of Life of Patients with Urolithiasis. J. Urol. 200, 1290–1294 (2018).

23. Harvey, J. A., Zobitz, M. M. & Pak, C. Y. Bioavailability of citrate from two different preparations of potassium citrate. J. Clin. Pharmacol. 29, 338–341 (1989).

24. Mazurek, M. P. et al. Molecular origin of plasma membrane citrate transporter in human prostate epithelial cells. EMBO reports 11, 431–437 (2010).

25. Rosenthal, A. K. et al. The progressive ankylosis gene product ANK regulates extracellular ATP levels in primary articular chondrocytes. Arthritis Res. Ther. 15, R154 (2013).

26. Costello, J. C. et al. Parallel regulation of extracellular ATP and inorganic pyrophosphate: roles of growth factors, transduction modulators, and ANK. Connect. Tissue Res. 52, 139–146 (2011).

27. Mitton-Fitzgerald, E., Gohr, C. M., Bettendorf, B. & Rosenthal, A. K. The Role of ANK in Calcium Pyrophosphate Deposition Disease. Curr. Rheumatol. Rep. 18, 25 (2016).

28. Albright, R. A. et al. ENPP1-Fc prevents mortality and vascular calcifications in rodent model of generalized arterial calcification of infancy. Nature Commun. 6, 10006 (2015).

29. Li, Q. et al. Mutant Enpp1asj mice as a model for generalized arterial calcification of infancy. Dis. Models Mech. 6, 1227–1235 (2013).

30. Russell, R. G. G. Bisphosphonates: From Bench to Bedside. Ann. N. Y. Acad. Sci. 1068, 367–401 (2006).

31. Dedinszki, D. et al. Oral administration of pyrophosphate inhibits connective tissue calcification. EMBO Mol. Med. 9, 1463–1470 (2017).

32. Borst, P., Váradi, A. & van de Wetering, K. PXE, a Mysterious Inborn Error Clarified. Trends Biochem. Sci. 44, 125–140 (2019).

33. Herrmann, M., Kinkeldey, A. & Jahnen-Dechent, W. Fetuin-A Function in Systemic Mineral Metabolism. TCM 22, 197–201 (2012).

34. Rüfenacht, H. S. & Fleisch, H. Measurement of inhibitors of calcium phosphate precipitation in plasma ultrafiltrate. Am. J. Physiol. 246, F648–55 (1984).

35. Engelke, U. F. H. et al. Guanidinoacetate methyltransferase (GAMT) deficiency diagnosed by proton NMR spectroscopy of body fluids. NMR Biomed. 22, 538–544 (2009).

36. Kirejczyk, J. K. et al. Urinary citrate excretion in healthy children depends on age and gender. Pediatr. Nephrol. 29, 1575–1582 (2014).

